# Genomic study of the Ket: a Paleo-Eskimo-related ethnic group with significant ancient North Eurasian ancestry

**DOI:** 10.1101/024554

**Authors:** Pavel Flegontov, Piya Changmai, Anastassiya Zidkova, Maria D. Logacheva, N. Ezgi Altimşik, Olga Flegontova, Mikhail S. Gelfand, Evgeny S. Gerasimov, Ekaterina E. Khrameeva, Olga P. Konovalova, Tatiana Neretina, Yuri V. Nikolsky, George Starostin, Vita V. Stepanova, Igor V. Travinsky, Martin Tříska, Petr Tříska, Tatiana V. Tatarinova

## Abstract

The Kets, an ethnic group in the Yenisei River basin, Russia, are considered the last nomadic hunter-gatherers of Siberia, and Ket language has no transparent affiliation with any language family. We investigated connections between the Kets and Siberian and North American populations, with emphasis on the Mal’ta and Paleo-Eskimo ancient genomes, using original data from 46 unrelated samples of Kets and 42 samples of their neighboring ethnic groups (Uralic-speaking Nganasans, Enets, and Selkups). We genotyped over 130,000 autosomal SNPs, identified mitochondrial and Y-chromosomal haplogroups, and performed high-coverage genome sequencing of two Ket individuals. We established that Nganasans, Kets, Selkups, and Yukaghirs form a cluster of populations most closely related to Paleo-Eskimos in Siberia (not considering indigenous populations of Chukotka and Kamchatka). Kets are closely related to modern Selkups and to some Bronze and Iron Age populations of the Altai region, with all these groups sharing a high degree of Mal’ta ancestry. Implications of these findings for the linguistic hypothesis uniting Ket and Na-Dene languages into a language macrofamily are discussed.

## Introduction

The Kets (an ethnic group in the Yenisei River basin, Russia) are among the least studied native Siberians. Ket language lacks apparent affiliation with any major language family, and is clearly distinct from surrounding Uralic, Turkic and Tungusic languages^1^. Moreover, until their forced settlement in 1930s, Kets were considered the last nomadic hunter-gatherers of North Asia outside the Pacific Rim^2^.

Ket language, albeit almost extinct, is the only language of the Yeniseian family that survived into the 21^st^ century. Most Yeniseian-speaking tribes (Arin, Assan, Baikot, Kott, Pumpokol, Yarin, Yastin) used to live south of the current Ket settlements. According to toponymic evidence, prior to the 17^th^ century speakers of this language family occupied vast territories of Western and Central Siberia, from northern Mongolia in the south to the middle Yenisei River in the north, and from the Irtysh River in the west to the Angara River in the east^3,4^. The Altai region was suggested as a homeland of the Yeniseian language family^2^, and ancestors of the Yeniseian people were tentatively associated^5^ with the Karasuk culture (3200-2700 YBP) of the upper Yenisei^6^. Yeniseian linguistic substrate is evident in many contemporary Turkic languages of this region (South Siberia): Altaian, Khakas, Shor, Tubalar, Tuvinian, and in Mongolic Buryat language^2^. As these languages are spoken in river basins with Yeniseian river names^1^, the Yeniseian tribes were likely to have mixed with these ethnic groups (and with the Southern Samoyedic groups Kamasins and Mators, now extinct^1^) at different times. We expect to find genetic signatures of these events.

Over the centuries, Kets and other Yeniseian people suffered relocation, extinction and loss of language and culture. First, they were under a constant pressure from the reindeer herders to the north (Enets and Nenets) and east (Evenks) and the Turkic-speaking pastoralists to the south. Second, Russian conquest of Siberia, which started at the end of the 16^th^ century, exposed the natives to new diseases, such as the 17^th^ century smallpox epidemic^7^. Third, in the 20^th^ century USSR resettled the Kets in Russian-style villages, thus interrupting their nomadic lifestyle^2^. Under the pressure of disease and conflict, the Kets have been gradually migrating north along the Yenisei River, and now reside in several villages in the Turukhansk district (Krasnoyarsk region); around 1,200 people in total^8^. Until the 20^th^ century, Kets, being nomadic hunters and fishers in a vast Siberian boreal forest, had little contact with other ethnic groups, which is manifested by the rarity of loanwords in Ket language^2^. However, since the collapse of the inter-Ket exogamous marriage system following the smallpox epidemics in the 17^th^ and 18^th^ centuries, Kets have been marrying Selkups, Uralic-speaking reindeer herders^2,9^. Moreover, during the 20^th^ century, the settled Kets have been increasingly mixing with other native Siberian people and with the Russians, which resulted in irrevocable loss of Ket language, genotype, and culture.

Recently, a tentative link was proposed between the Yeniseian language family and the Na-Dene family of Northwest North America (composed of Tlingit, Eyak, and numerous Athabaskan languages), thus forming a Dene-Yeniseian macrofamily^10–12^. The Dene-Yeniseian-linkage is viewed by some as the first relatively reliable trans-Beringian language connection^11^, with important implications for timing of the alleged Dene-Yeniseian population split, the direction of the subsequent migration (from or to America), the possible language shifts and population admixture^13–15^.

So far, no large-scale population study was conducted with samples from each of the presently occupied Ket villages. Previously, six Ket individuals were genotyped^16–18^ and two of them sequenced^19^ These studies concluded that the Kets do not differ from surrounding Siberian populations, a rather surprising finding, given their unique language and ancient hunter-gatherer life-style. In order to clarify this issue, in 2013 and 2014 we collected 57 (46 unrelated) samples of Kets and 42 unrelated samples of their neighboring Uralic-speaking ethnic groups (Nganasans inhabiting the Taymyr Peninsula, and Enets and Selkups living further south along Yenisei). We genotyped approximately 130,000 autosomal SNPs and determined mitochondrial and Y-chromosomal haplogroups with the GenoChip array^20^. We also performed high-coverage genome sequencing of two Ket individuals. Using these data, we investigated connections between Kets and modern and ancient Siberian and North American populations (including the Mal’ta and Saqqaq ancient genomes). In addition, we estimated Neanderthal contribution in Kets’ genome and in specific gene groups.

Mal’ta is a ~24,000 YBP old Siberian genome, recently described^21^ as a representative of ancient North Eurasians (ANE)^22^, a previously unknown northeastern branch of the Eurasian Paleolithic population. ANE contributed roughly 30-40% to the gene pool of Native Americans of the first settlement wave^21^ and about 50% to the Bronze Age Yamnaya culture in the Pontic-Caspian steppe^6,23,24^. Massive expansion of the Corded Ware culture around 5,000-4,000 YBP, originating from the Yamnaya source, introduced the ANE genetic pool into Central and Western Europe and thus reshaped its genetic landscape^6,23^. During the same period, the Afanasievo and Andronovo cultures, genetically similar to the Yamnaya culture, expanded into the Altai region (South Siberia) and later mixed with Siberian populations, giving rise to the Bronze Age Karasuk culture and later Iron Age cultures^6^.

A global maximum of ANE ancestry occurs in Native Americans, with lower levels in peoples of more recent Beringian origin, i.e. indigenous populations of Chukotka, Kamchatka, the Aleutian Islands and the American Arctic^21,22,25^. In modern Europe, ANE genetic contribution is the highest in the Baltic region, on the East European Plain and in the North Caucasus^6,22,23^. However, little is known about the distribution of ANE ancestry in its Siberian homeland. According to a single *f_4_* statistic, the Kets had the third highest value of ANE genetic contribution among all Siberian ethnic groups, preceded only by Chukchi and Koryaks^26^. Thus, we suggest that the Kets might represent the peak of ANE ancestry in Siberia; the hypothesis we tested extensively in this study. We also investigated continuity between the modern Kets and Altaians, and the ancient Bronze and Iron Age populations of the Altai region discussed above: the Karasuk culture samples dated to 3531-3261 YBP, and Iron Age samples roughly dated to 2900-1100 YBP^6^.

The 4,000 YBP genome from Greenland^16^ represents the Saqqaq archeological culture (4,500-2,800 YBP). This culture formed a continuum with Dorset and Norton cultures (2,500-1,000 YBP), collectively termed Paleo-Eskimo^25^. Paleo-Eskimos were culturally and genetically distinct from modern Inuits and Eskimos^13,25^. The Saqqaq culture is a part of the wider Arctic Small Tool tradition (ASTt) that had rapidly spread across Beringia and the American Arctic coastal regions (but not the interior) after 4,800 YBP, bringing pottery, bow and arrow to the northern North America^13,15,27^. According to the archaeological data, the likely source of this spread was located in Siberia, namely in the Lena River basin (probably, in the Bel’kachi culture^13^). On genetic grounds, Paleo-Eskimos were also argued to represent a separate migration into America^16,25,28^. ASTt spread coincided with the arrival of mitochondrial haplogroup D2 into America and the spread of haplogroup D2a^29^; the Saqqaq individual bore haplogroup D2a1^30^. The closest modern relatives of Saqqaq occur among Beringian populations (Chukchi, Koryaks, Inuits^25^) and Siberian Nganasans^16^. In addition, Saqqaq has been linked to Na-Dene-speaking Chipewyans (16% contribution to this population modeled with admixture graphs^28^). However, mitochondrial haplogroup data^25,30,31^ argues against the proximity of Paleo-Eskimos to contemporary Na-Dene people^13,15^, primarily due to the very high frequency of haplogroup A in the latter^32^. Archeological evidence seems to support this argument^13^.

There is no archaeological evidence of considerable trans-Beringian population movements between the inundation of the Bering Platform around 13,000-11,000 YBP and 4,800 YBP. Therefore, it is unlikely that the hypothetical Dene-Yeniseian language family has separated prior to 11,000 YBP, according to current concepts of time depth in language evolution^13,15^, and hence ASTt could be the vehicle spreading Dene-Yeniseian languages and genes from Siberia to Alaska and to the American Arctic^13^. However, as argued based on language phylogenetic trees^33^ in the framework of the Beringian standstill model^29,34^, the Dene-Yeniseian languages have originated in Beringia and spread in both directions. Irrespective of the migration direction and their relationship to contemporary Na-Dene groups, Paleo-Eskimos are the primary target for investigating genetic relationship with the Kets.

In this study, we found that: (1) Kets and Selkups constitute a clade closely related to Nganasans; (2) Nganasans, Kets, Selkups, and Yukaghirs form a cluster of populations most closely related to Paleo-Eskimos in Siberia (not considering indigenous populations of Chukotka and Kamchatka); (3) unlike Nganasans, Kets derive roughly 30-40% of their ancestry from ancient North Eurasians; (4) Kets show genetic continuity with the hypothetical homeland of Yeniseian languages, as they are closely related to the ancient individuals of the Karasuk culture and to the later Iron Age individuals from the Altai.

## Results and Discussion

### Identification of a non-admixed Ket genotype

We compared the GenoChip SNP array data for the Ket, Selkup, Nganasan, and Enets populations (Suppl. file S1) to the worldwide collection of populations^35^ based on 130K ancestry-informative markers^20^.We applied GPS^35^ and reAdmix^36^ algorithms to infer provenance of the samples and confirm self-reported ethnic origin (see details in Suppl. Information, Section 4, Suppl. Figs. 4.2, 4.3, Suppl. Table 3). Combining the two algorithms, we identified a subset of non-admixed Kets among self-identified Ket individuals, and nominated two individuals for whole-genome sequencing. ADMIXTURE^37^ revealed very similar profiles of components in individuals sequenced in this study and in two (Fig. 1A) or four (Suppl. Fig. 5.1) Kets from published sources. We note that proportion of the European component at K=4 was slightly lower in the published Ket individuals (28% vs. 32% on average).

**Fig. 1.**
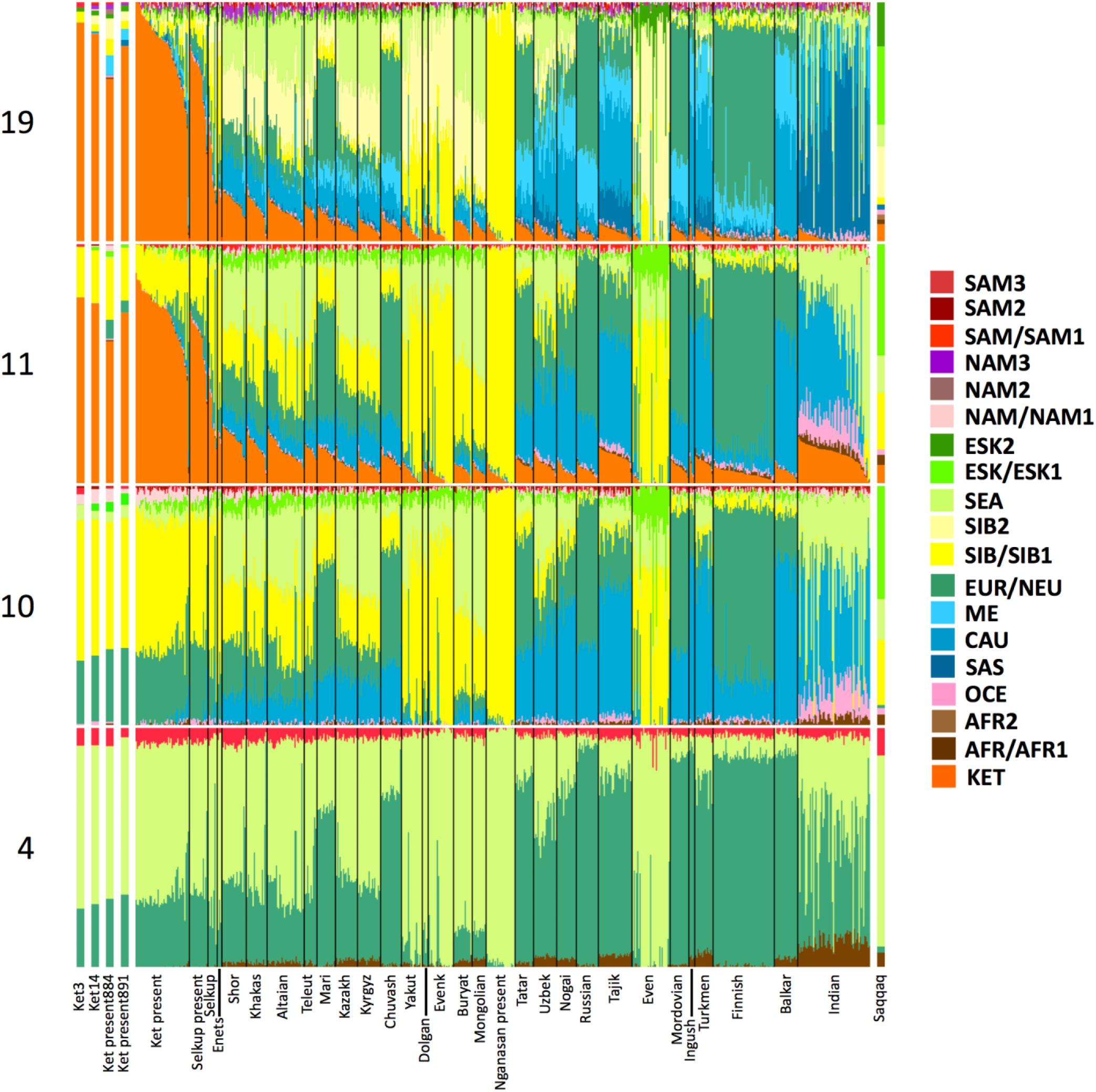
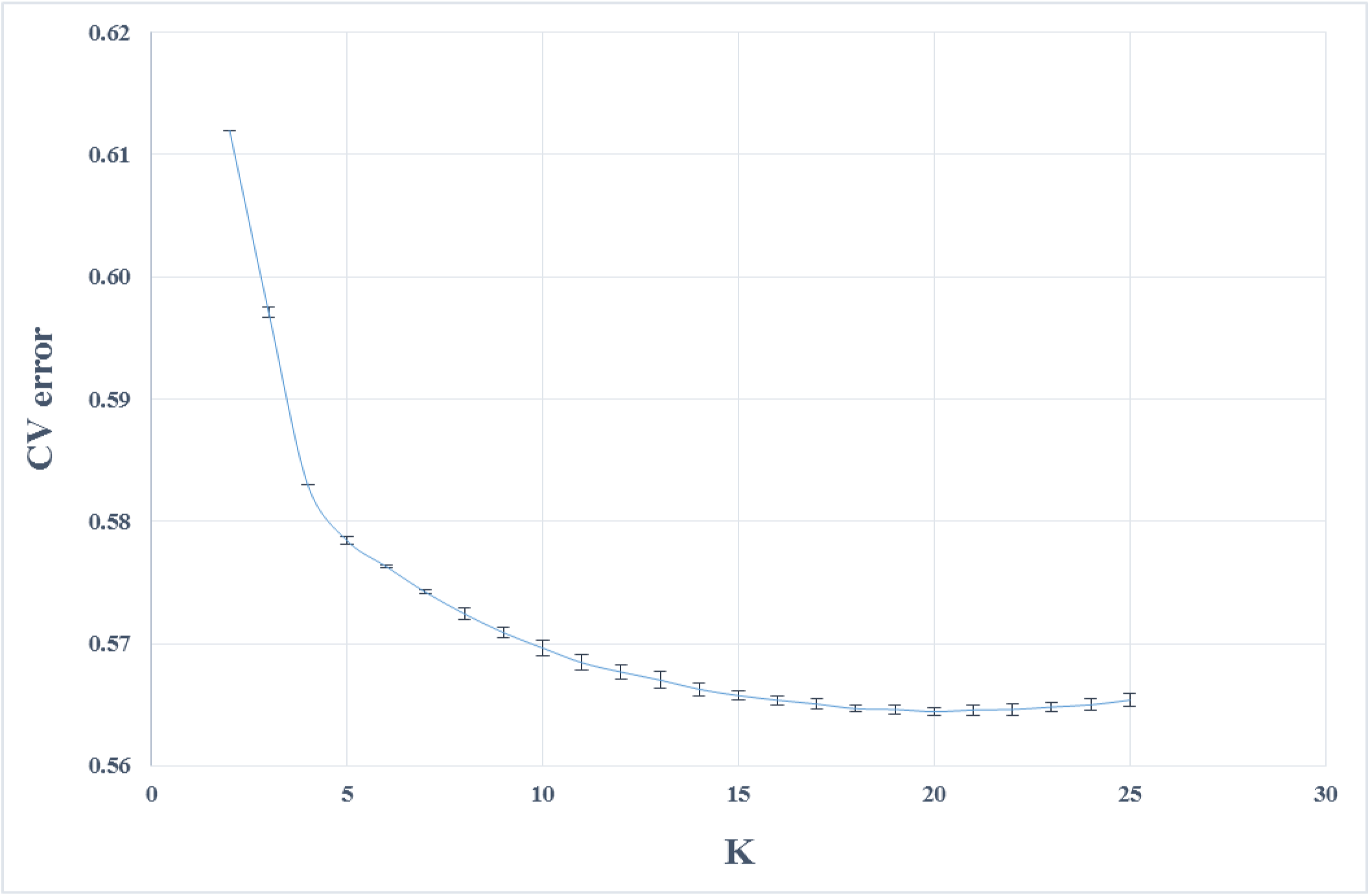
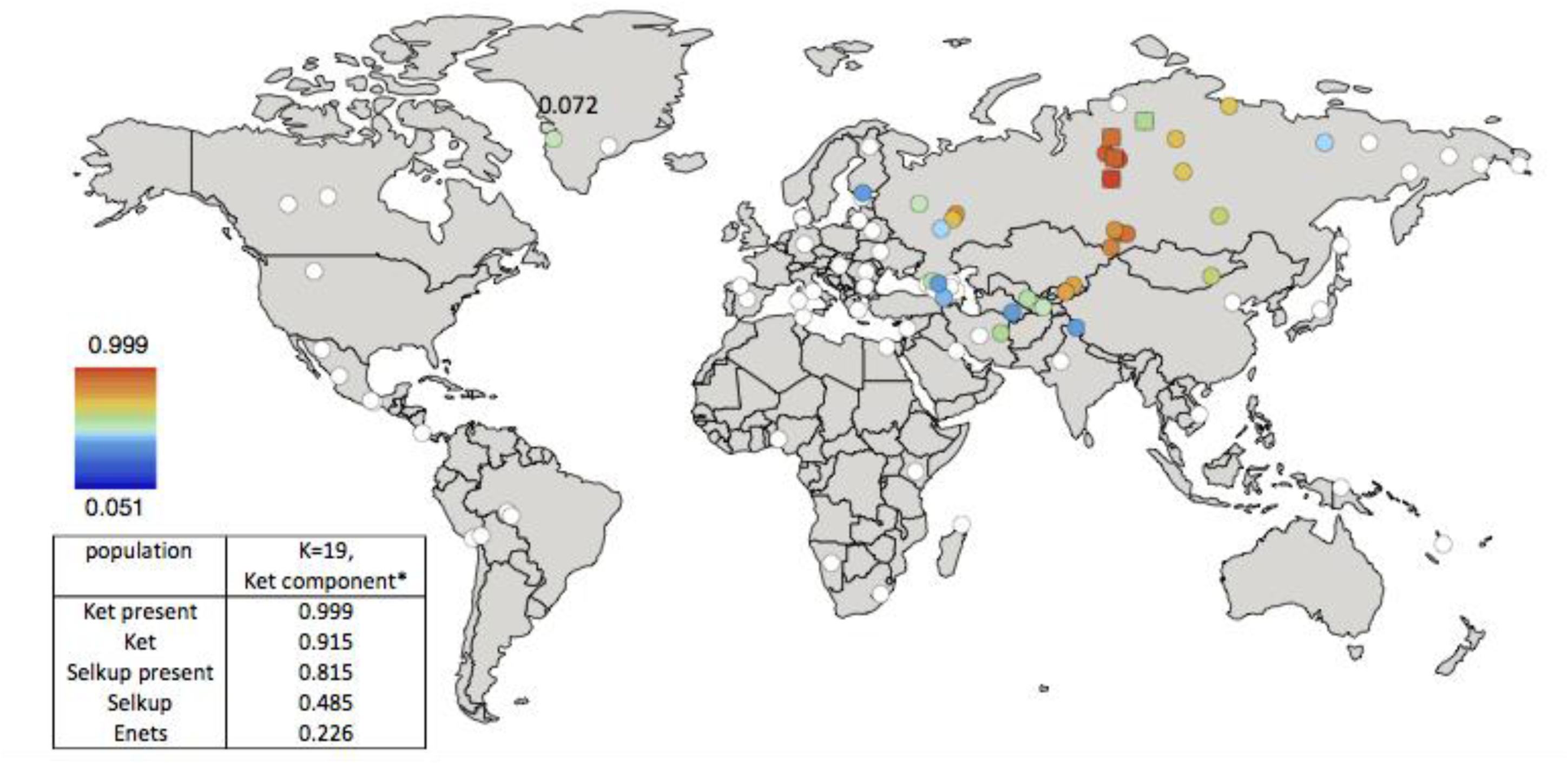
**A**. Admixture coefficients plotted for dataset ‘GenoChip + Illumina arrays’. Abbreviated names of admixture components are shown on the left as follows: SAM, South American; NAM, North American; ESK, Eskimo (Beringian); SEA, South-East Asian; SIB, Siberian; NEU, North European; ME, Middle Eastern; CAU, Caucasian; SAS, South Asian; OCE, Oceanian; AFR, African. The Ket-Uralic (‘Ket’) admixture component appears at K≥11, and admixture coefficients are plotted for K=4, 10, 11, and 19. Although K=20 demonstrates the lowest average cross-validation error, the Ket-Uralic component splits in two at this K value, therefore K=19 was chosen for the final analysis. Only populations containing at least one individual with >5% of the Ket-Uralic component at K=19 are plotted, and individuals are sorted according to values of the Ket-Uralic component. Admixture coefficients for the Saqqaq ancient genome are shown separately on the right, and for two reference Kets and two Ket individuals from this study – on the left. **B**. Average cross-validation (CV) error graph with standard deviations plotted. Ten-fold cross-validation was performed. The graph has a minimum at K=20. **C**. Color-coded values of the Ket-Uralic admixture component at K=19 plotted on the world map using QGIS v.2.8. Maximum values in each population are taken, and only values >5% are plotted. Top five values of the component are shown in the bottom left corner, and the value for Saqqaq is shown on the map.

### ‘Ket-Uralic’admixture component

Using the GenoChip SNP array^20^, we genotyped 130K ancestry-informative markers in the Ket, Selkup, Nganasan, and Enets populations (Suppl. file S1). Following the exclusion of first-, second-, and third-degree relatives among the individuals genotyped in this study (Suppl. file S1, Suppl. Fig. 4.1), we merged the GenoChip array data with the published SNP array datasets to produce a worldwide dataset of 90 populations and 1,624 individuals, focused on Siberia and America (Suppl. Table 2). The intersection dataset, containing 32,189 SNPs (Suppl. Table 1), was analyzed with ADMIXTURE^37^ (Fig. 1). At K≥11, ADMIXTURE identified a characteristic component for the Ket population (Suppl. Information Section 5). This component reached its global maximum of nearly 100% in Kets, closely followed by Selkups from this study (up to 81.5% at K=19), the reference Selkups (up to 48.5%) and the Enets (up to 22.6%). The difference between the Selkups from this study and the reference Selkups^21^ can be attributed to a much closer geographic proximity of the former population to the settlements of Kets, with whom they have a long history of cohabitation and mixture^2,10^.

The ‘Ket’ component occurred at high levels (up to ~20%) in four Turkic-speaking populations of the Altai region: Shors, Khakases, Altaians, and Teleuts. Notably, the Altai region was populated by Yeniseian-speaking people before they were forced to retreat north (Suppl. Information, Section 2). Lower levels of the ‘Ket’ component, from 5% to 15%, were observed in the following geographic regions (in decreasing order): the Volga-Ural region, Central and South Asia, East Siberia and Mongolia, and North Caucasus. The ‘Ket’ component also occurred at a low level in Russians (up to 7.1%), Finns (up to 5.4%), and, remarkably, in the Saqqaq ancient genome from Greenland (7.2%, see below).

In order to verify and explain the geographic distribution of the ‘Ket’ admixture component, we have performed ADMIXTURE analysis on two additional datasets, different in population (Suppl. Table 2) and marker selection (Suppl. Table 1) (see Suppl. Information, Section 5). In summary, we suggest the existence of an admixture component with a peculiar geographic distribution, observed in some previous studies but not discussed there^17,18^. In addition to the Kets, this component is characteristic also for Samoyedic-speaking and Ugric-speaking people of the Uralic language family: Selkups, Enets, Nenets, Khanty, Mansi, with a notable exception of Samoyedic-speaking Nganasans. The proportion of the ‘Ket-Uralic’ admixture component correlated strongly with the worldwide frequency of mitochondrial haplogroup U4 (Pearson’s correlation coefficient up to 0.8 and a corresponding *p*-value of 7×10^−8^) and with the frequency of Y-chromosomal haplogroup Q in Eurasian populations (correlation coefficient up to 0.9 and *p*-value 2×10^−7^) (Suppl. Information, Section 10).

### Kets in the context of Siberian populations

In order to study the relationship of Kets and other Siberian populations with the relevant ancient genomes, we have constructed three additional datasets: the dataset based on the Ket genome sequences and the HumanOrigins array SNP data^22^, and two datasets based on genome sequences only (Suppl. Tables 1 and 2). The Ket and Selkup populations were closely related according to multiple analyses (see the ADMIXTURE plot in Fig. 1, PCA plots in Suppl. Figs. 6.3, 6.6, TreeMix tree in Fig. 2, and outgroup *f_3_* statistics^38^ in Suppl. Fig. 7.2). Nganasans appeared as the closest relatives of Kets according to statistics *f_3_*(Yoruba; Ket, X): the statistic for Nganasans was significantly different from that of the second-best hit (Suppl. Fig. 7.2). In general, outgroup *f_3_* statistics (Yoruba; Test, X) were tightly correlated between the Kets, the Selkups, and the Nganasans, with Pearson’s correlation coefficients ranging from 0.96 to 0.999 (Suppl. Information, Section 7), suggesting that these populations form a closely related group. In line with these results, Nganasans, Kets, Selkups, and Yukaghirs formed a clade in a maximum likelihood tree constructed with TreeMix on a HumanOrigins-based dataset of 194,750 SNPs (Fig. 2).

**Fig. 2.**
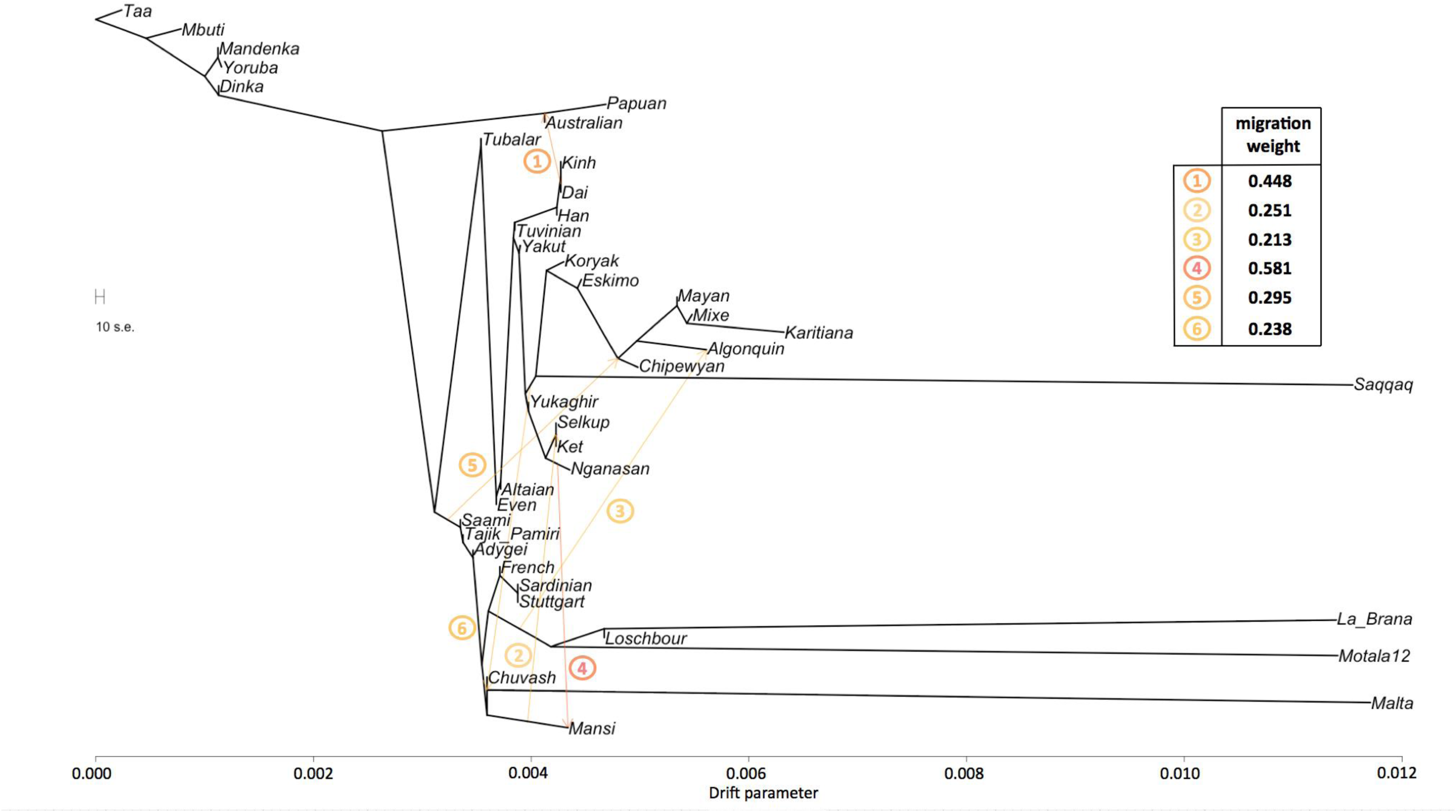
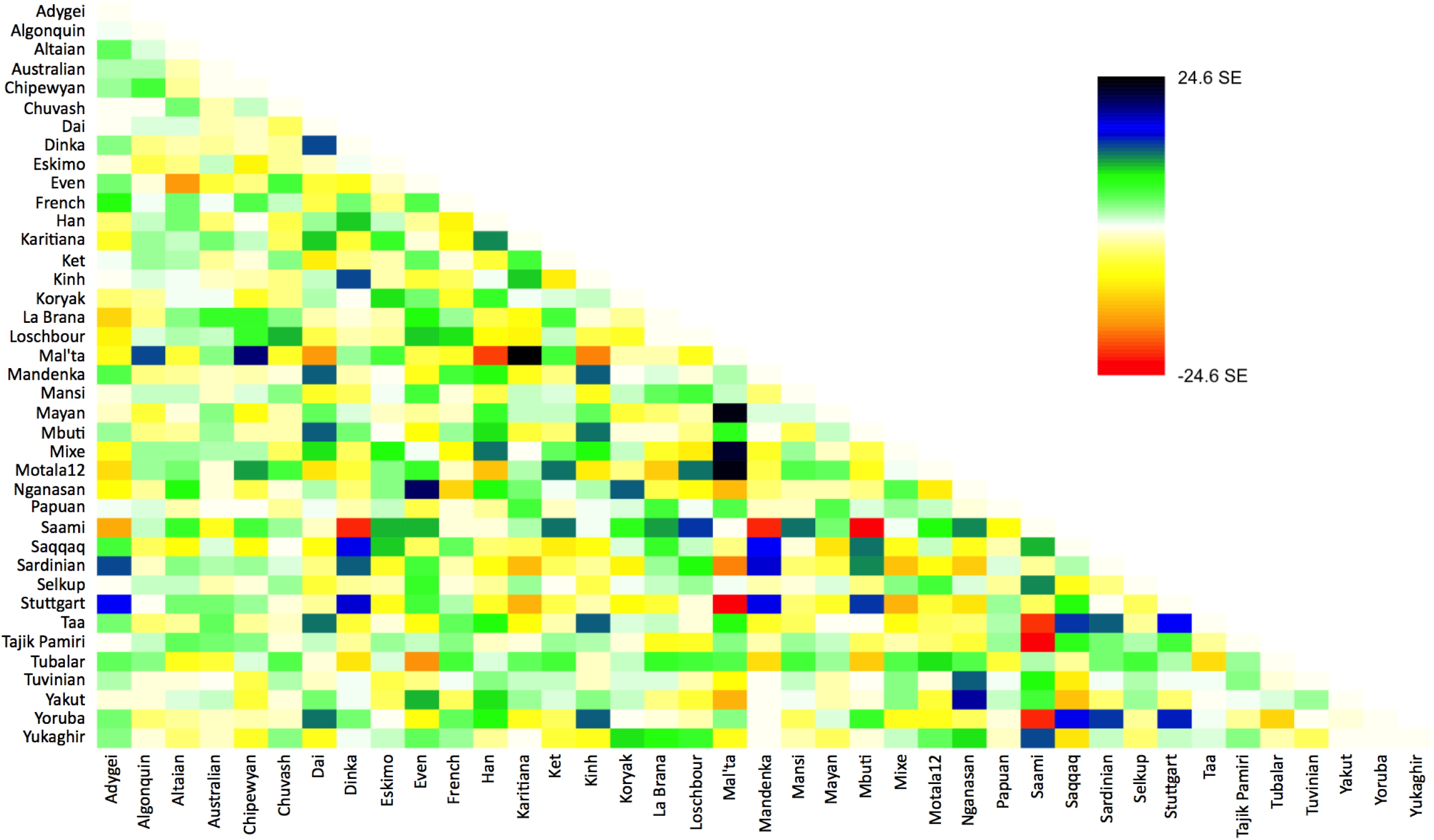
**A**. A maximum likelihood tree with 6 migration edges computed on the dataset ‘Ket genomes + HumanOrigins’ with selected populations (194,750 SNPs, 39 populations, 527 individuals). Drift parameter is shown on the x-axis. **B**. Residuals from the fit of the model to the data visualized. 98% of variance is explained by the tree.

In our ADMIXTURE analyses (Fig. 1A, Suppl. Fig. 5.4), the Saqqaq Paleo-Eskimo individual featured the following components: Beringian, Siberian, and South-East Asian. Thus, Saqqaq Paleo-Eskimo has mostly Beringian ancestry (similar to modern Eskimo, Inuits, Aleutians, Koryaks, etc.): see outgroup *f_3_* statistics and associated Z_diff_ scores in Suppl. Figs. 7.17-7.19, migration edges modelled with TreeMix in Fig. 3, and the ADMIXTURE results in the original study^16^. Beringian ancestry in Saqqaq is combined with considerable Siberian ancestry: 32% or 28% as a sum of Siberian ADMIXTURE components in this study (Fig. 1A, Suppl. Fig. 5.4); ~25% according to ADMIXTURE analysis in the original study (Rasmussen et al. 2010)^16^; from 31% to 57% according to *f_4_* statistic ratios calculated with various outgroups (Suppl. Information, Section 8, Suppl. Table 6). This ‘core Siberian’ component in Saqqaq is apparently most closely related to modern Nganasans^16^ (Suppl. Fig. 7.17) and to the Nganasan-related clade in general (see a TreeMix tree in Fig. 2). The Kets are the only representatives of this clade in the genome-based datasets in this study. According to the pairwise correlation between outgroup *f_3_* statistics (the method used in Allentoft et al.^6^), Kets are closer to Saqqaq as compared to Nivkhs, Altaians, Buryats, and Yakuts (Suppl. file S2). According to Euclidean distances in the ten-dimensional space of principal components on the HumanOrigins dataset, Kets were a closer population to Saqqaq than Nganasans, Selkups, Yukaghirs, and the other populations (Fig. 5). However, the outgroup *f_3_* statistics (Yoruba; Saqqaq, Ket) in many cases were not significantly different from *f_3_*(Yoruba; Saqqaq, other Siberian population): e.g., see |Z_diff_| scores < 3 for *f_3_*(Yoruba; Saqqaq, Nivkh) (Suppl. Figs. 7.18, 7.19). The same result was produced with *f_4_*(Saqqaq, Yoruba; Nivkh, Ket): an absolute Z-score was lower than 2 (Suppl. Fig. 8.18A,B).

**Fig. 3.**
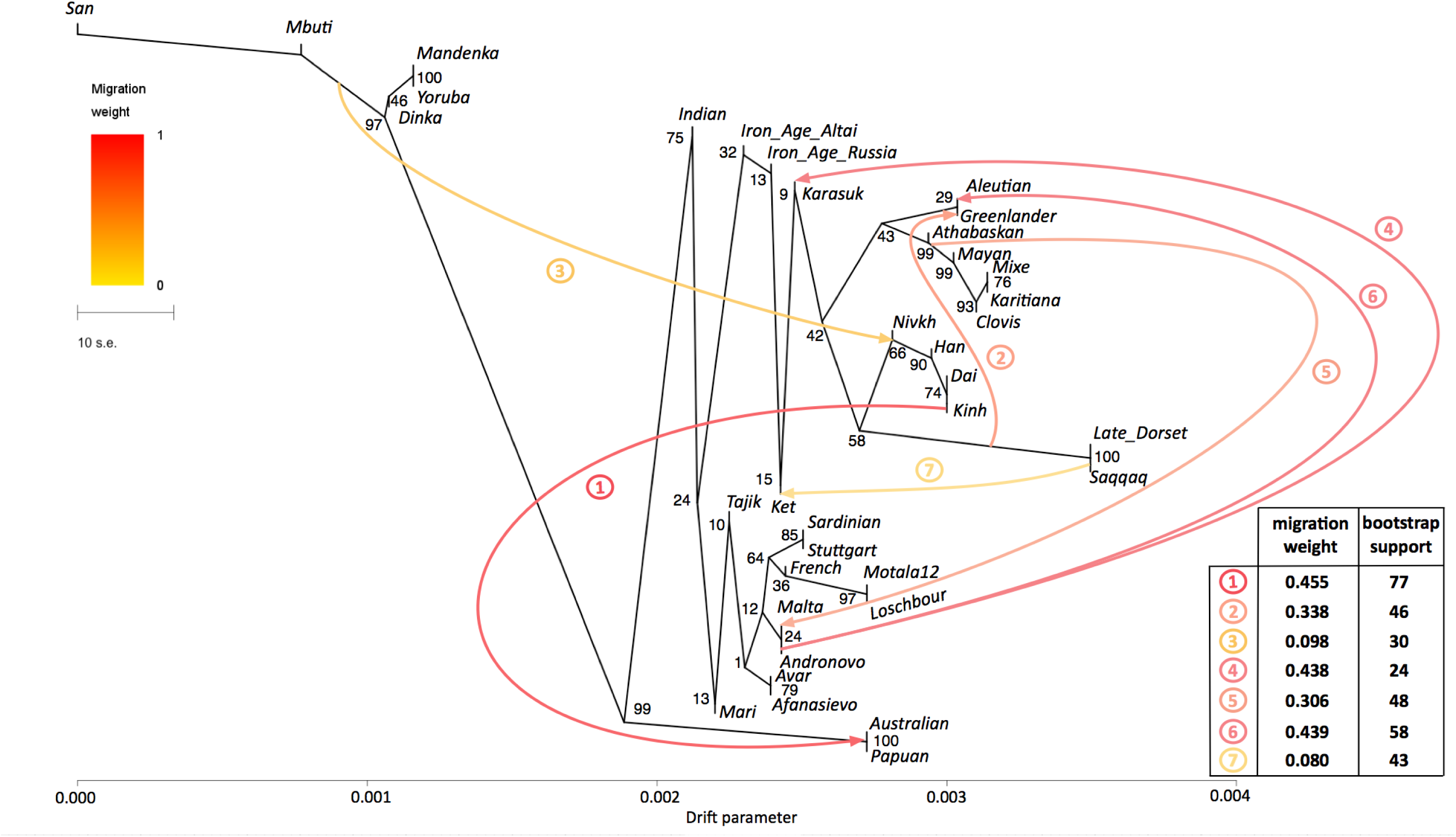
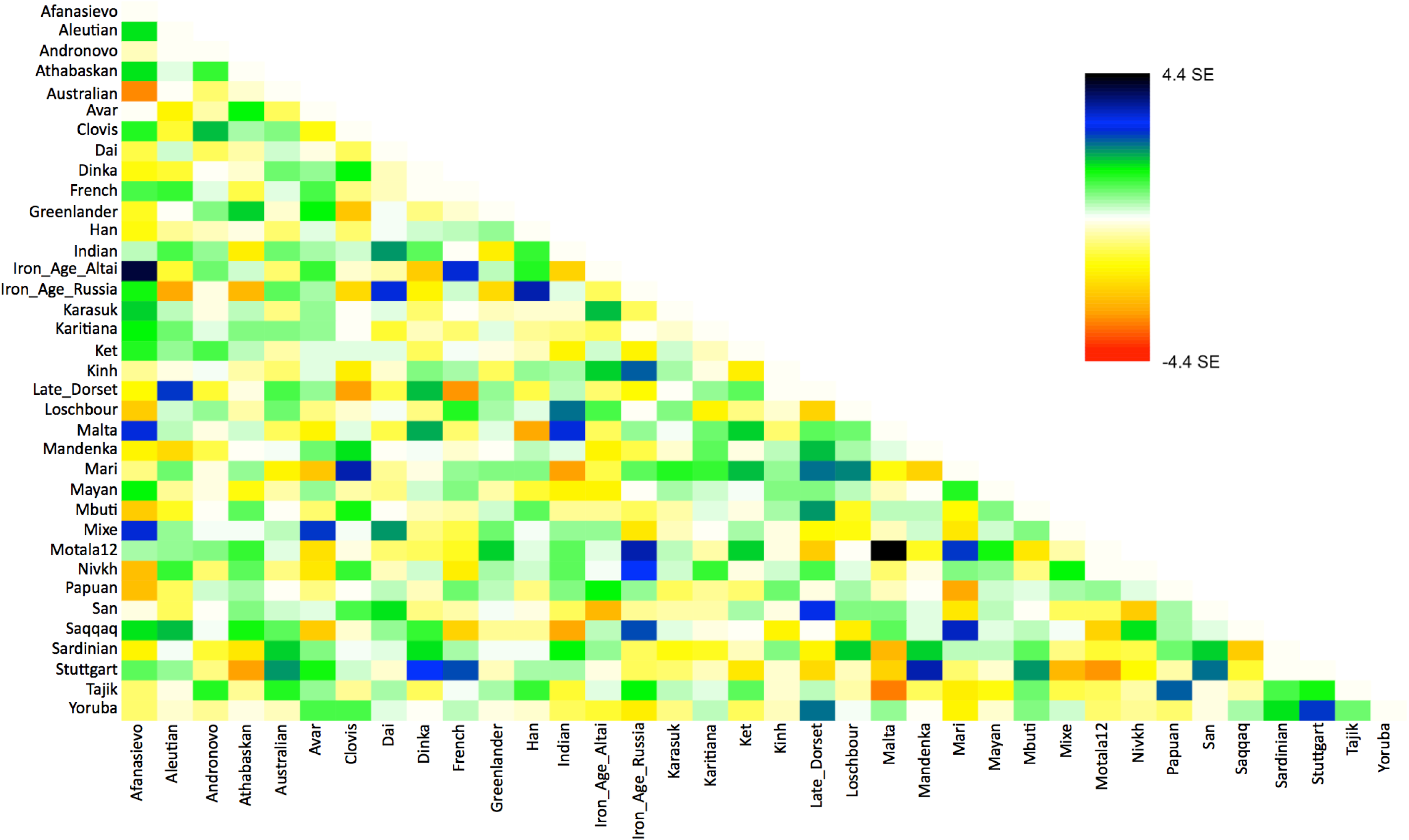
**A**. A maximum likelihood tree with 7 migration edges computed on the genome-based dataset without transitions. Edge weight and bootstrap support values are shown in the table, the drift parameter is shown on the x-axis, and bootstrap support values for tree nodes are indicated. Migration edges are numbered according to the order of their appearance in the sequence of trees from *m*=0 to *m*=8. Note to the figure: as migration edges and tree topology are inter-dependent in bootstrapped trees, bootstrap support for the edges in the original tree was calculated by summing up support for closely similar edges in bootstrapped trees. Below these edge groups are listed for edges #1-7: 1/ Australian and/or Papuan ⇔ the (Nivkh, Han, Dai, Kinh) clade or any of its members; 2/ Greenlander Inuit or the (Greenlander, Aleutian) clade ⇔ Saqqaq and/or Late Dorset (optionally a wider clade with Nivkh); 3/ any clade containing African populations ⇔ any clade composed of Nivkh/Han/Dai/Kinh (optionally a wider clade with Late Dorset and/or Saqqaq and/or Iron Age Altai); 4/ any clade composed of Mal’ta/Afanasievo/Andronovo (optionally a wider clade with Aleutian and/or Mari) ⇔ Karasuk; 5/ Mal’ta (optionally a wider clade with Motala12/Afanasievo/Andronovo/Aleutian) ⇔ any clade composed exclusively of Native Americans and/or Greenlander; 6/ any clade composed exclusively of populations with European ancestry ⇔ Aleutian; 7/ Ket (optionally a wider clade with Karasuk and/or Iron Age Altai and/or Iron Age Russia) ⇔ Saqqaq and/or Late Dorset. **B**. Residuals from the fit of the model to the data visualized. 96.72% variance is explained by the tree.

Allentoft et al.^6^ have shown that in the early Bronze Age the Altai region was inhabited by a genetically West Eurasian population of the Afanasievo archaeological culture, most similar to the Yamnaya culture with about 50% of ANE ancestry^6^, and to the modern Avars (Suppl. file S2, Fig. 3, Suppl. Figs. 9.4, 9.5). In the late Bronze Age this population apparently had being gradually admixed with Siberians, giving rise to the Karasuk culture and the later cultures of the Iron Age^6^. The ancient genome ‘Iron Age Russia’, carbon-dated to 721-889 AD^6^, is most closely related to the typical modern Siberians: Nganasans according to the outgroup *f_3_* statistic in the original study^6^ and Altaians or Koryaks according to the outgroup *f_3_* statistics and their pairwise correlations on our datasets lacking Nganasans (Suppl. file S2). However, according to various analyses, the ‘Iron Age Altai’ (dated roughly to 2900-1500 YBP) and the Karasuk (carbon-dated to 3531-3261 YBP) populations of two and six genomes, respectively, are most closely related to each other and to Kets (Suppl. files S2, S3, Suppl. Figs. 7.5, 7.6, 8.17A, 9.1). The outgroup *f3* statistics (Yoruba; Karasuk, X) on both genome-based datasets selected Kets as the best hit for Karasuk (Suppl. files S2, S3, Suppl. Fig. 7.5), although statistics for Mayans, Greenlanders, Mixe, Saqqaq, Mal’ta, Iron Age Russia, and Aleutian were not significantly different (|Z_diff_| score < 3). Similarly, Native American, Beringian populations and Selkups were the best hits for Iron Age Altai and Karasuk according to the outgroup *f_3_* statistics in the original study (Kets were lacking in the dataset, see Allentoft, et al.^6^). Importantly, the Karasuk culture has been tentatively associated with the Yeniseian-speaking people based on the toponymic evidence^5^, and the Altai region is considered to be the homeland of the Yeniseian language family^2^. As another piece to this puzzle, we observed genetic continuity between the Kets and the ancient genomes from the Altai.

### Mal’ta (ancient North Eurasian) ancestry in Kets

The outgroup statistic *f_3_*(Yoruba; Mal’ta, Ket) (Raghavan et al. 2014)^21^ was higher than statistics for all other Siberian and most Beringian populations. However, the statistic values were not significantly different within a large group of North Eurasians, according to Z_diff_ scores (Suppl. Figs. 7.13-7.16). Z_diff_ score for *f_3_*(Yoruba; Mal’ta, Ket) vs. *f_3_*(Yoruba; Mal’ta, Nganasan) equaled 7.4, 7.0 vs. *f_3_*(Yoruba; Mal’ta, Yukaghir), and only 2.7 vs. *f_3_*(Yoruba; Mal’ta, Selkup) (Suppl. Fig. 7.13). Thus, we suggest that, unlike the other members of the Nganasan-related clade (Fig. 2), Kets and, to a lesser extent, Selkups have a high proportion of Mal’ta ancestry, alternatively referred to as the ANE ancestry^22^. The Mal’ta ancestry in Kets was further supported by the TreeMix^39^ analysis, specifically by a migration edge connecting Mal’ta to the Ket-Karasuk clade, with a weight of 43% (Suppl. Fig. 9.1, Suppl. Table 7). Taking into account the admixture coefficients for the two sequenced Ket individuals (Ket891 and Ket884, Fig. 1A), we selected Ket891 as an individual with lower values of the North European and Siberian admixture components (in the K=19 dimensional space). In addition, Ket891 was identified as non-admixed by reAdmix analyses (Suppl. Table 3). Ket891 demonstrated a slightly closer genetic affinity to Mal’ta (compare Suppl. Figs. 7.13 and 7.14).

These results were consistent with calculations of *f_4_* statistic in two configurations: (X, Chimp; Mal’ta, Stuttgart) or (X, Papuan; Sardinian, Mal’ta), reproducing the previously used statistics^22,26^ (Suppl. Figs. 8.1-8.6). *f_4_*(X, Chimp; Mal’ta, Stuttgart) analysis tests whether the population X has more drift shared with Mal’ta or with Stuttgart (an early European farmer, EEF^22^). Sardinians were used as the closest modern proxy for EEF^22^ in *f_4_*(X, Papuan; Sardinian, Mal’ta). All possible population pairs (X,Y) were tested by *f_4_*(Mal’ta, Yoruba; Y, X) on the genome-based dataset, including both Ket individuals (Fig. 4A). Compared to Kets, Mal’ta was significantly closer to none of the populations including Native Americans.

**Fig. 4.**
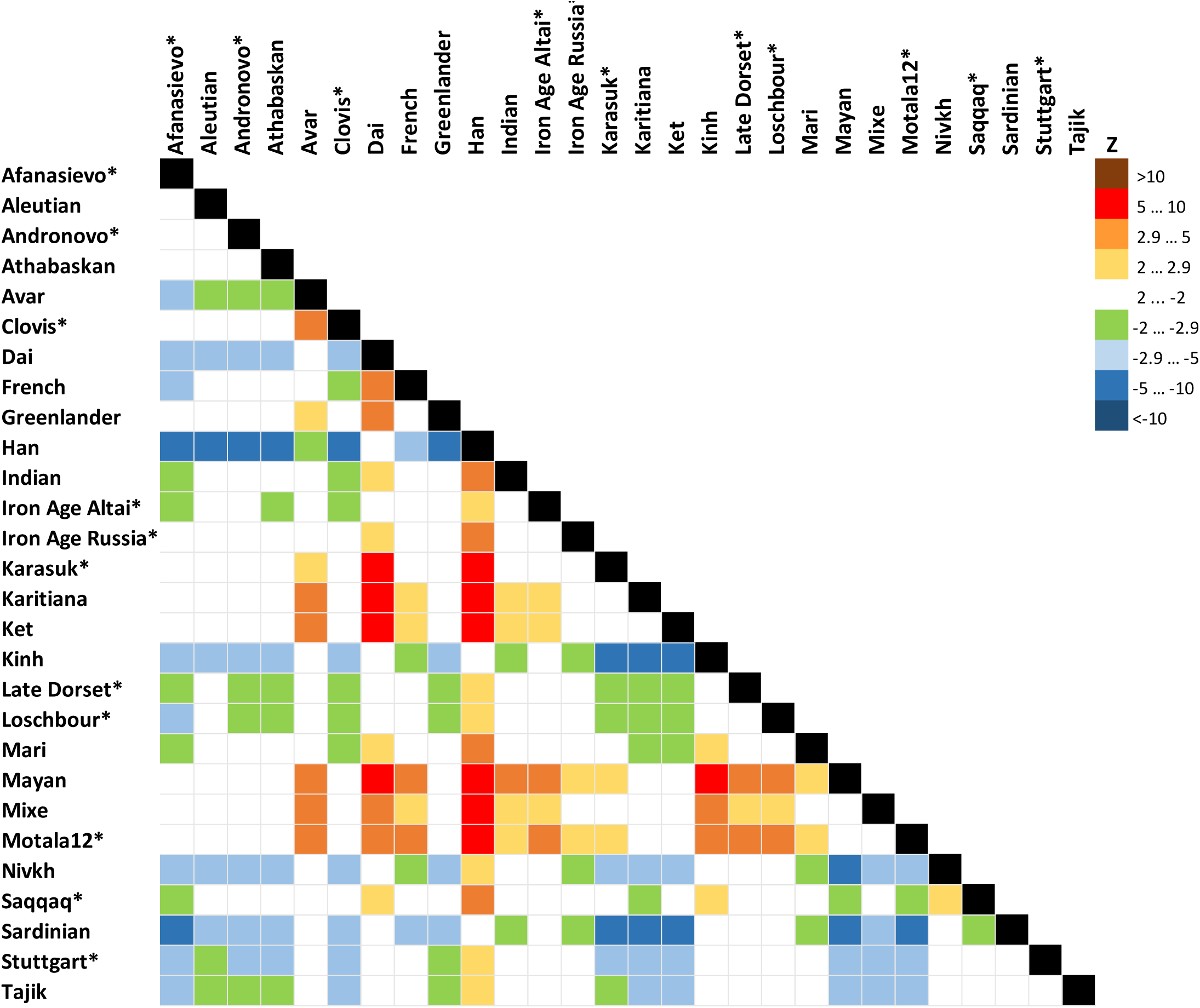
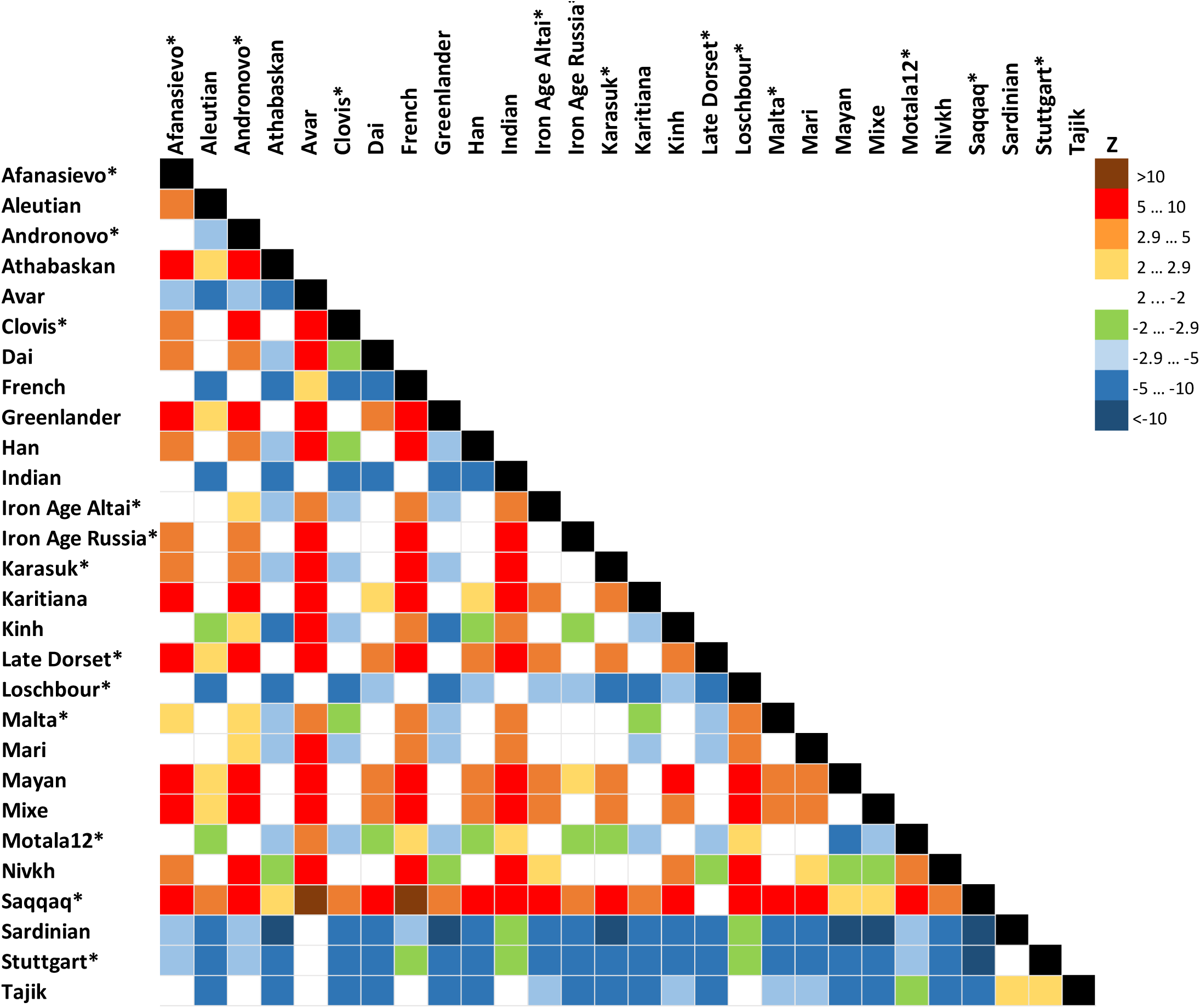
Statistics *f_4_*(Mal’ta, Yoruba; Y, X) (**A**), *f_4_*(Ket, Yoruba; Y, X) (**B**) computed on the genome-based dataset with African, Australian and Papuan populations excluded. See the corresponding results for the dataset without transitions in Suppl. Figs. 8.15 and 8.16, respectively. A matrix of color-coded Z-scores is shown, and ancient genomes are marked with asterisks. Z-score equals the number of standard errors by which the statistic differs from zero, and |Z| > 2.9 demonstrates that the statistic is significantly different from zero using Bonferroni correction for 27 independent tests (threshold p-value of 0.001852). Rows show Z-scores for *f_4_*(Mal’ta, Yoruba; row, column) or *f_4_*(Ket, Yoruba; row, column), *vice versa* for columns.

**Fig. 5.**
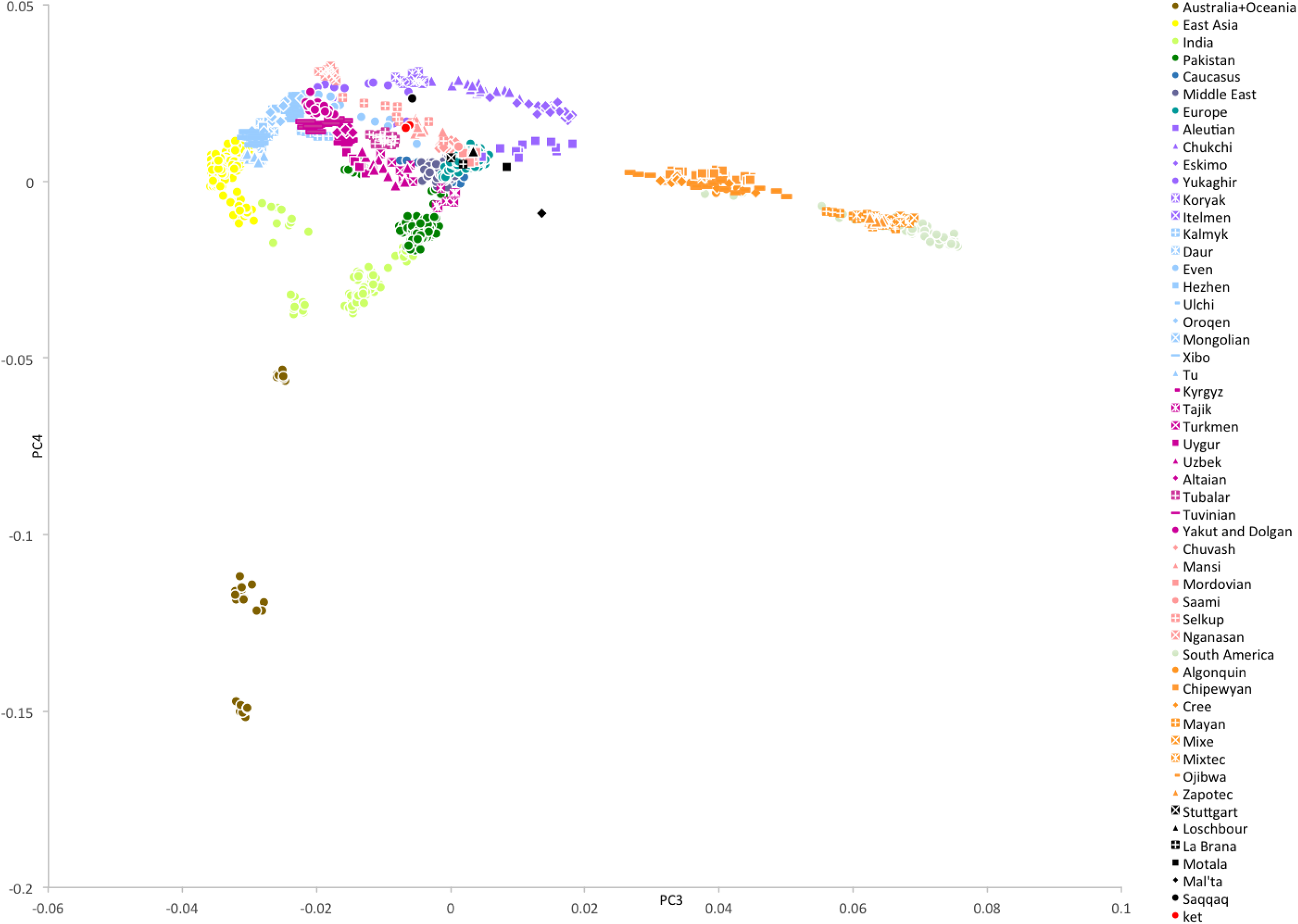
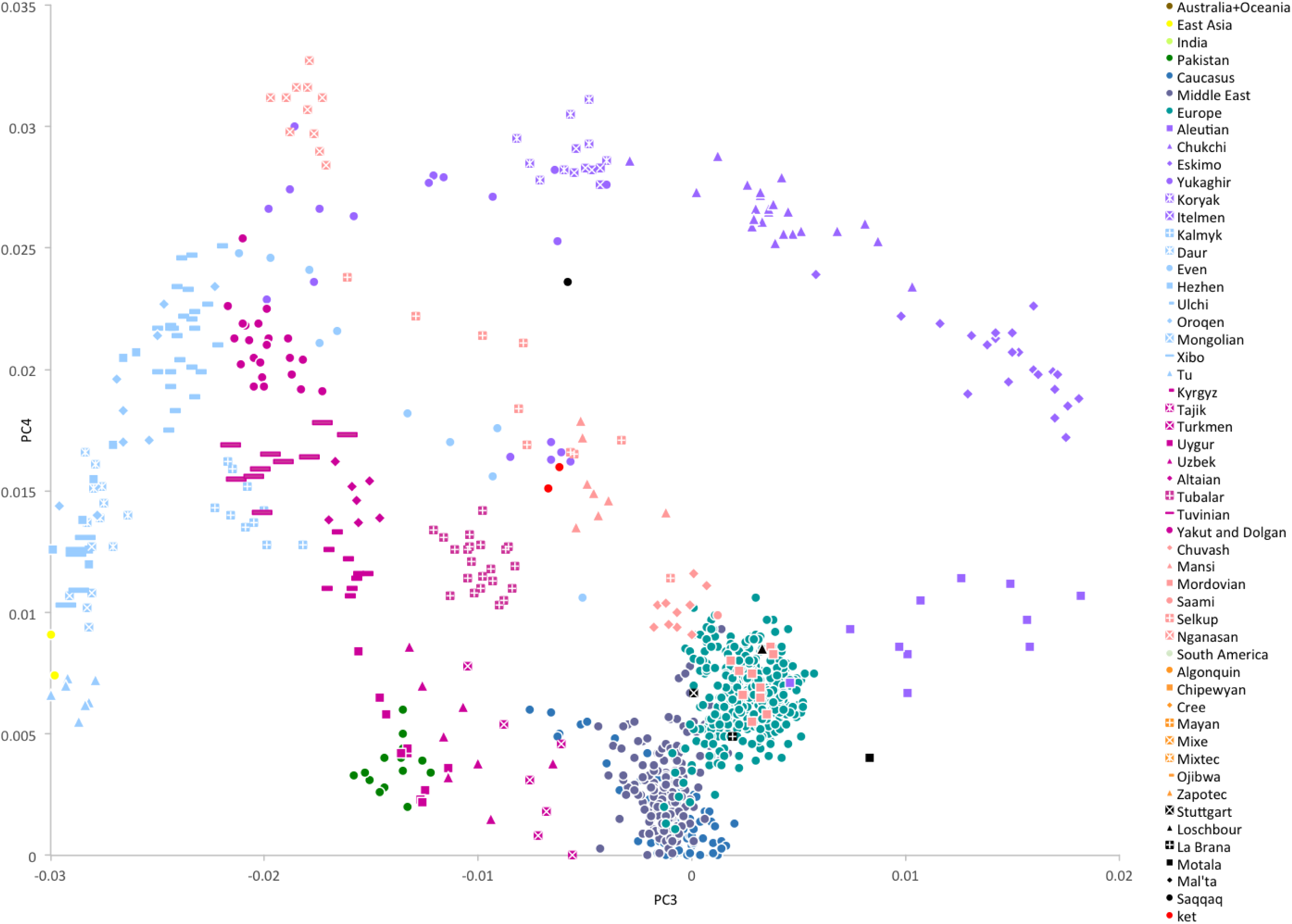
**A**. PC3 vs. PC4 plot for the dataset ‘Ket genomes + HumanOrigins array’. African populations are not shown. Populations are color-coded by geographic region or language affiliation (in the case of Siberian and Central Asian populations), and most relevant populations are differentiated by marker shapes. Ancient genomes are shown in black. For the corresponding PC1 vs. PC2 plot see Suppl. Fig. 6.7. **B**. PC3 vs. PC4 plot, zoom on the Ket individuals. Here is a list of populations closest to Saqqaq based on the average Euclidean distances in the multi-dimensional space of ten principal components (distances in parentheses): Ket (0.022), Nganasan (0.025), Selkup (0.026), Yukaghir (0.028), Eskimo (0.032), Koryak (0.032), Mansi (0.032), Itelmen (0.033), Chukchi (0.033), Dolgan (0.035).

Based on all analyses, we can tentatively model Kets as a two-way mixture of broadly defined East Asians and ANE. Therefore, ANE ancestry in Kets can be estimated, using various *f_4_*-ratios, at 27% to 43% (depending on reference populations and datasets), vs. 25 ‒ 53% in various Native American groups (Suppl. Table 6, see details in Suppl. Information, Section 8). Given non-significant Z_diff_ scores and *f_4_* statistics (Fig. 4A) discussed above, it is difficult to identify the exact Eurasian population west of Chukotka and Kamchatka with the highest degree of the Mal’ta ancestry, but Kets are a good candidate. We speculate that ANE component was acquired by ancestors of Kets in the Altai region, where the Bronze Age Okunevo culture was located, with a surprisingly close genetic proximity to Mal’ta^6^. Later, the Yeniseian-speaking people occupied this region until the 16^th^-18^th^ centuries^3,4^.

### Kets and Na-Dene speakers

In this study, Na-Dene-speaking people were represented by Athabaskans, Chipewyans, Tlingit, and, possibly, Haida. The latter language was originally included into the Na-Dene language family^40^, although this affiliation is now disputed^10^ Na-Dene-speaking people were suggested to be related, at least linguistically, to Yeniseian-speaking Kets^10^. ADMIXTURE, PCA, *f_3_* and *f_4_* statistics, and TreeMix analyses failed to identify a link between Kets and Athabaskans, Chipewyans, or Tlingit (see Suppl. Information, Sections 7 and 8). TreeMix constructed trees where Athabaskans or Chipewyans formed a stable highly supported clade with other Native Americans (e.g., bootstrap support 99, Fig. 3A). This topology was supported by statistics *f_4_*(Athabaskan, Yoruba; Ket, X), which demonstrated significantly negative Z-scores < -5, for Clovis, Greenlanders, Karitiana, Mayans, and Mixe (Suppl. Fig. 8.19) on the genome-based dataset with or without transitions. Notably, the same topology was previously demonstrated for Athabaskans^19^.

Notably, the Arctic Small Tool tradition the Saqqaq culture belongs to, may reflect the Dene-Yeniseian movement over the Bering Strait^13,15^. According to the admixture graph analysis modeling relationships among Chipewyans, Saqqaq, Algonquin, Karitiana, Zapotec, Han, and Yoruba^28^, only one topology fits these data. In this topology, the Chipewyans represent a mixture of 84% First Americans and 16% Saqqaq^28^. Our estimates using *f_4_*-ratios are roughly similar: 4-15% Saqqaq ancestry in Chipewyans and 0-9% in Athabaskans (Suppl. Table 6). Considering 57% as the highest proportion of Siberian ancestry in Saqqaq obtained in this study, we predict up to ~9.1% of Siberian ancestry in Chipewyans, i.e. 57%×16% of Saqqaq ancestry in Chipewyans. Similarly, only 1.2% (noise level) of the Ket-Uralic admixture component is predicted in Chipewyans, with 7.2% as the highest percentage of this component found in Saqqaq (Fig. 1C). Given such low levels of expected genetic signal, we cannot reliably test the hypothetical genetic connection between Yeniseian and Na-Dene-speaking people, provided the employed methods and population samples. Moreover, considerable Beringian ancestry in Saqqaq makes the Saqqaq ancestry in Chipewyans difficult to distinguish from potential admixture with Eskimo and Inuits, representatives of the third settlement wave^25,28^. Hopefully, the question of the Dene-Yeniseian genetic relationship and its correlation with the linguistic relationship will be answered with a study of autosomal haplotypes and/or very rare allelic variants in relevant genomic data, which would test whether a relatively recent Siberian (Nganasan- and Ket-related) gene flow occurred into Chipewyans, for example, and whether it was mediated by Paleo-Eskimos.

## Conclusions

Based on previous studies^16,25,28^, the Saqqaq individual, and the Paleo-Eskimos in general^25^ may represent a separate and relatively recent migration into America. The Paleo-Eskimos have large proportions of Beringian (i.e. Chukotko-Kamchatkan and Eskimo-Aleut), Siberian, and South-East Asian ancestry. We have also shown that Kets and Selkups belong to a group of modern populations closest to an ancient source of Siberian ancestry in Saqqaq. This group also includes, but is probably not restricted to, Uralic-speaking Nganasans and Yukaghirs (the latter speak an isolated language). Unlike the other populations of this group, Kets, and, to a lesser degree Selkups, have a high proportion of Mal’ta (ancient North Eurasian) ancestry.

As shown previously^28^, Chipewyans, a modern Na-Dene-speaking population, have about 16% of Saqqaq ancestry. Thus, a gene flow dated at 5,000-6,000 YBP^16^ can be traced from the cluster of Siberian populations to Saqqaq, and from Saqqaq to Na-Dene. However, the genetic signal in contemporary Na-Dene-speaking ethnic groups is substantially diluted. The genetic proximity of Kets to the source of Siberian ancestry in Saqqaq correlates with the hypothesis that Na-Dene languages of North America are specifically related to Yeniseian languages of Siberia, now represented by Ket language only^10^. However, this genetic link is indirect and requires further study of population movement and language shifts in Siberia.

In addition, we show that Kets represent a modern Siberian population closest to ancient individuals of the Karasuk culture, spanning a period from about 3400 YBP to 2900 YBP (the individuals analyzed were dated to 3531-3261 YBP), and to few investigated Iron Age individuals of the Altai region (2900-1100 YBP)^6^. This genetic continuity correlates with historical linguistic data suggesting that the homeland of Yeniseian languages was located in the Altai region^2,5^.

## Methods

### Sample Collection

Saliva samples were collected and stored in the lysis buffer (50 mM Tris, 50 mM EDTA, 50 mM sucrose, 100 mM NaCl, 1% SDS, pH 8.0) according to the protocol of Quinque et al.^41^ The following cities and villages along the Yenisei River were visited (Suppl. Fig. 1.1): Dudinka (69.4°, 86.183°), Ust’-Avam (71.114°, 92.821°), Volochanka (70.976°, 94.542°), Potapovo (68.681°, 86.279°), Farkovo (65.720°, 86.976°), Turukhansk (administrative center of the Turukhanskiy district, 65.862°, 87.924°), Baklanikha (64.445°, 87.548°), Maduika (66.651°, 88.428°), Verkhneimbatsk (63.157°, 87.966°), Kellog (62.489°, 86.279°), Bakhta (68.841°, 96.144°), Bor (61.601°, 90.018°), Sulomai (61.613°, 91.180°). Volunteers rinsed their mouths with cold boiled water, and then collected up to 2 mL of saliva into a tube filled with 3 mL of the buffer. Samples were stored at environmental temperature (ranging from 4°C to 30°C) for up to two weeks, and DNA was isolated within one month after sample collection. Information about ethnicity, place of birth, and about first-, second-, and third-degree relatives was provided by the volunteers. The study was approved by the ethical committee of the Lomonosov Moscow State University (Russia), and the study methods were carried out in accordance with the approved guidelines and regulations. All volunteers have signed informed consent forms. The study was approved by local administrations of the Taymyr and Turukhansk districts and discussed with local committees of small Siberian nations for observance of their rights and traditions.

### DNA extraction

DNA extraction protocol was adapted from the high-salt DNA extraction method^41^. DNA extracted from saliva represents a mix of human and bacterial DNA, and their ratio was checked by quantitative (q) PCR with 2 primer pairs (human: 1e1 5’-GTCCTCAGCGCTGCAGACTCCTGAC-3’, BG1R 5’-CTTCCGCATCTCCTTCTCAG-3’; bacterial: 8F 5’-AGAGTTTGATCCTGGCTCAG-3’, 519R 5’-GWATTACCGCGGKGCTG-3’). The PCR reaction mixture included: 13,2 μL of sterile water, 5 μL of qPCR master mix PK154S (Evrogen, Moscow, Russia), 0,4 μL of each primer and 1 μL of DNA sample. Amplification was performed with thermocycler StepOnePlus Applied Biosystems™ (Life Technologies, USA) with the standard program. Most DNA samples had low levels of bacterial contamination, and were used for further analysis: 22% of samples had the human/bacterial DNA ratio <1; 59% of samples had the ratio from 1 to 2, 19% of samples had the ratio >2.

### Genotyping

GenoChip (the Genographic Project’s genotyping array)^20^ was used for genotyping 158 individuals (see details in Suppl. file S1). GenoChip includes ancestry-informative markers obtained for modern populations, the ancient Saqqaq genome, and two archaic hominins (Neanderthal and Denisovan), and was designed to identify all known Y-chromosome and mitochondrial haplogroups. The chip allows genotyping about 12,000 Y-chromosomal and approximately 3,300 mitochondrial SNPs, and over 130,000 autosomal and X-chromosomal SNPs. Genotyping was performed at the GenebyGene sequencing facility (TX, USA).

### Control of Sex Assignment

In order to avoid mix-ups in sex assignment, we compared heterozygosity of X chromosome and missing rate among Y-chromosomal SNPs across the samples. All female samples had >62% Y-chromosomal SNPs missing and X chromosome heterozygosity >0.138, while male samples demonstrated values <1.2% and <0.007, respectively. Four wrong sex assignments were corrected based on these thresholds.

### Genome sequencing and genotype calling

Genome sequencing has been performed for Ket individuals 884 (a male born in Baklanikha, mitochondrial haplogroup H, Y-chromosomal haplogroup Q1a2a1) and 891 (a female born in Surgutikha, mitochondrial haplogroup U5a1d). Prior to genome sequencing, we used NEBNext Microbiome DNA Enrichment Kit (New England Biolabs, USA) in order to enrich the samples for human DNA. This kit exploits the difference in methylation between eukaryotes and prokaryotes through selective binding of CpG-methylated (eukaryotic) DNA by the MBD2 protein. We took 500 ng of DNA from each sample and processed it according to manufacturer’s instructions. Success of the enrichment was estimated using qPCR with primers for human RPL30 and bacterial 16S rRNA genes. The resulting human-enriched fraction was used for library construction using the TruSeq DNA sample preparation kit (Illumina, USA). We made libraries also from non-enriched DNA in order to assess whether enrichment leads to biases in sequence coverage. Libraries from enriched and non-enriched DNA were sequenced using the HiSeq2000 instrument (Illumina, USA) with read length 101+101 bp, two lanes for each library. As both enriched and non-enriched libraries produced similar coverage profiles and similar SNP counts in test runs of the *bcbio-nextgen* genotype calling pipeline (data not shown), their reads were pooled for subsequent analyses. Resulting read libraries for samples 884 and 891 had a median insert size of 215 bp and 343 bp, and coverage of 61x and 44x, respectively.

The *bcbio-nextgen* pipeline v. 0.7.9 (https://bcbio-nextgen.readthedocs.org/en/latest/) has been used with default settings for the whole read processing workflow: adapter trimming, quality filtering, read mapping on the reference genome hg19 with BWA, duplicate read removal with Picard, local realignment, SNP calling and recalibration with GATK v3.2-2, and annotation against dbSNP_138 with snpEff. Two alternative genotype calling modes have been tested in GATK: batch genotype calling for several samples, emitting only sites with at least one non-reference allele in at least one individual (GATK options -- standard_min_confidence_threshold_for_calling 30.0 -- standard_min_confidence_threshold_for_emitting 30.0, -- emitRefConfidence at default); or calling genotypes for each sample separately, emitting all sites passing the coverage and quality filters (GATK options --standard_min_confidence_threshold_for_calling 30 -- standard_min_confidence_threshold_for_emitting 30 --emitRefConfidence GVCF -- variant_index_type LINEAR --variant_index_parameter 128000). The former approach maximized the output of dbSNP-annotated sites (7.36-7.62 million sites per individual vs. ~3.7 million sites for the latter approach), and therefore was used to generate calls subsequently merged with various SNP array and genomic datasets (Suppl. Table 1). To increase sensitivity of this approach, genotype calling has been performed with GATK HaplotypeCaller for six genomes in one run: Kets 884 and 891, two Yoruba, and two Kinh (Vietnamese) individuals downloaded from the 1000 Genome Project database^42^. We chose Yoruba samples NA19238 and NA19239, and Vietnamese samples HG01873 and HG02522 as they had read coverage similar to the Ket samples.

### Combined Datasets

Combined datasets generated in this study and their properties are listed in Suppl. Table 1. In all cases, datasets were designed to maximize population coverage in Russia (Siberia in particular), in Central Asia and in the Americas, while keeping only few reference populations in the Middle East, in South and South-East Asia, Africa, Australia, and Oceania. Third-degree and closer relatives detected through questionnaires and pedigree analysis and individuals of mixed ethnicity were excluded from the Enets, Ket, Nganasan, and Selkup population samples. Overall, 88 of 158 individuals remained (see supposed percentage of relatedness for each sample pair and a list of selected samples in Suppl. file S1 and hierarchical clustering of all samples based on genetic distance in Suppl. Fig. 4.1). All datasets underwent filtering using PLINK^43^ v. 1.9. Maximum missing rate per SNP thresholds of 0.03 or 0.05 were used (Suppl. Table 1), except for dataset ‘Ket genomes + reference genomes’, for which a more relaxed threshold of 0.11 was used to accommodate the Mari individual and ancient genomes with low coverage^21^ and to keep the high number of SNPs at the same time. Linkage disequilibrium (LD) filtering was applied to all datasets except for the GenoChip-based one since SNPs included into the GenoChip array underwent LD filtering with the r^2^ threshold of 0.4 as described in Elhaik et al.^20^. For the other datasets the following LD filtering settings were used: window size of 50 SNPs, window step of 5 SNPs, r^2^ threshold 0.5 (PLINK v. 1.9 option ‘--indep-pairwise 50 5 0.5’). Ancient genome data within the HumanOrigins dataset were provided as artificially haploid: one allele was selected randomly at each diploid site, since confident diploid calls were not possible for low-coverage ancient genomes (see further description of the approach in Lazaridis et al.^22^). We applied the same procedure to ancient genomes included into dataset ‘Ket genomes + reference genomes’. Other details pertaining to individual datasets are listed below and in Suppl. Table 1, and their population composition is shown in Suppl. Table 2.

**GenoChip + Illumina arrays** (alternative name ‘GenoChip-based’). The dataset was constructed by merging selected populations genotyped with GenoChip^35^, genotyping data obtained with GenoChip in this study, and SNP array data (various Illumina models) from the following sources: Behar et al.^44^; Cardona et al.^45^; Fedorova et al.^17^; Li et al.^46^; Kidd et al.^47^; Raghavan et al.^21^; Rasmussen et al.^16^; Reich et al.^28^; Silva-Zolezzi et al.^48^; Surakka et al.^49^; Yunusbayev et al.^50^. Three ancient genomes, La Braña^51^, Saqqaq^16^, and Clovis^52^, were also added (genotypes in VCF format were obtained from the respective publications). Only SNPs included into the GenoChip array were used and further filtered as described above. Maximum missing rate per individual was 50%, but three ancient genomes (Clovis, Saqqaq and La Braña) were exempt. After filtering, the dataset contained 1,624 individuals from 90 populations and 32,189 SNPs, and included published data for two Ket individuals^17^.

**Ket genomes + Illumina arrays**. In order to include populations relevant for our analyses, e.g. Burusho, Khanty, and Nenets, omitted from the previous dataset due to very low marker overlaps, full-genome SNP calls for two Ket individuals (see above) were merged with SNP array data (various Illumina models) from the following sources: HapMap3^53^; Behar et al.^44,54^; Cardona et al.^45^; Fedorova et al.^17^; Li et al.^46^; Raghavan et al.^21^; Rasmussen et al.^16^; Reich et al.^28^; Silva-Zolezzi et al.^48^; Yunusbayev et al.^50^. The filtered dataset contained modern individuals only: 2,549 individuals from 105 populations and 103,495 SNPs, and had low missing SNP rates (maximum missing rate per individual 0.04). The dataset included published data for four Ket individuals^16,17^, and was used for ADMIXTURE analysis only.

**Ket genomes + HumanOrigins array** (alternative name ‘HumanOrigins-based’). For analyzing relevant ancient genomes alongside the Ket genomes in context of multiple modern populations genotyped with the HumanOrigins Affymetrix SNP array^22,38^, the full dataset of Lazaridis et al.^22^ was merged with Ket genome data and filtered (LD filtering with the r^2^ threshold of 0.5, maximum per SNP missing rate of 0.05). The resulting set contained 217 populations/genomes. In order to make the dataset more manageable computationally and more focused, 78 populations/genomes were removed, keeping the following ancient genomes with coverage >1x: Neanderthal and Denisovan genomes; La Braña 1^51^, Loschbour^22^, Mal’ta/MA1^21^, and Motala12^22^ (Eurasian hunter-gatherers); Stuttgart (a west European farmer of the LBK archaeological culture^22^) and Saqqaq^16^. The final dataset contained 1,786 individuals from 139 populations and 195,918 SNPs.

**Ket genomes + HumanOrigins array + Verdu et al. 2014.** The final version of the previous dataset ‘Ket genomes + HumanOrigins array’ was merged with Illumina 610-Quad SNP array genotyping data for six North American native populations (Haida, Nisga’a, Splatsin, Stswecem’c, Tlingit, Tsimshian^55^) and filtered (Suppl. Table 1). The major reason for constructing this dataset was the inclusion of Tlingit and Haida, Na-Dene-speaking populations not present in the other datasets, and the dataset was used only for analyses focused on Na-Dene-speaking populations. The final dataset contained 1,867 individuals from 145 populations and 68,625 SNPs.

**Ket genomes + reference genomes** (alternative name ‘genome-based’). The following 19 ancient genomes were included into the dataset: Clovis^52^, Late Dorset^25^ and Saqqaq^16^ Paleo-Eskimos, Mal’ta (an ANE representative^21^), Motala12 and Loschbour (WHG^22^), Stuttgart (EEF^22^), and 12 ancient genomes with coverage >1x were selected from archaeological cultures of the Altai region (labelled according to the original publication as Afanasievo, Andronovo, Iron Age Altai, Iron Age Russia, and Karasuk)^6^. To ensure dataset uniformity, genotype calling for these ancient genomes was performed *de novo* in a batch run, instead of using published genotypes generated with different genotype calling protocols. Ancient DNA reads mapped on the reference genome hg19 (provided by their respective authors) were used for genotype calling with the ANGSD software v. 0.800^56^ with the following settings: SAMtools calling mode (option -GL 1); genotype likelihood output (option -doGlf 2); major allele specified according to the reference genome (-doMajorMinor 4); allele frequency obtained based on the genotype likelihoods (-doMaf 1); SNP *p*-value 10^−6^. The resulting genotype likelihood files were transformed into genotypes in the VCF format using BEAGLE utilities gprobs2beagle and beagle2vcf, with a minimum genotype likelihood cut-off of 0.6. Subsequently one allele was selected randomly at each diploid site, since confident diploid calls were not possible for low-coverage ancient genomes. Genotype data for ancient genomes were merged with the following modern samples using PLINK v. 1.9 (and subsequently filtered as described in Suppl. Table 1):(i) 7.36-7.62 million GATK genotype calls for two Kets, two Yoruba, and two Vietnamese individuals; and (ii) one Aleutian, two Athabaskans, two Greenlanders, two Nivkhs^25^; one Avar, one Indian, one Mari, one Tajik^21^; one Australian aboriginal^57^, one Karitiana, one Mayan^52^; Simons Genome Diversity Project panels A^58^ and B^59^ containing 25 genomes of 13 populations. The final dataset contained 64 individuals from 36 populations and 398,163 SNPs.

**Ket genomes + reference genomes without transitions**. In order to mitigate the effect of ancient DNA deamination and the resulting C to T substitutions^60^, we constructed another version of the genome-based dataset, with all CT and AG SNPs excluded prior to the LD filtering step. The final dataset contained 64 individuals from 36 populations and 189,964 SNPs.

**Ket genomes + Raghavan et al. 2015 with/without transitions**. In order to include two additional Ket genomes, genomes of Altaians, Buryats, Yakuts, Koryaks, and Eskimo (ten in total), and three genomes of Native Americans (Huichol, Mayan, Tsimshian) published by Raghavan et al.^19^, we merged these data with the genome-based dataset described above prior to linkage disequilibrium filtering and transition polymorphisms removal. Upon filtering, the resulting datasets contained 79 individuals from 43 populations, 225,010 SNPs in the original version, and 104,727 SNPs with transitions excluded. These genome-based datasets were helpful due to an extended sampling of Siberian populations, but because of a relatively small number of markers they were used for few selected analyses only (Suppl. Table 1).

### PCA

The principal component analysis (PCA) was carried out in the *smartpca* program included in the EIGENSOFT package^61^. We calculated 10 eigenvectors and ran the analysis without removing outliers. Then we computed the Euclidean distance in the space of ten principal components between all pairs of individuals, and calculated mean distance between the Saqqaq individual and all modern populations.

### ADMIXTURE analysis

The ADMIXTURE software implements a model-based Bayesian approach that uses block-relaxation algorithm in order to compute a matrix of ancestral population fractions in each individual (Q) and infer allele frequencies for each ancestral population (P)^37^. A given dataset is usually modeled using various numbers of ancestral populations (K). For each K from 2 to 25, 100 analysis iterations were generated with different random seeds. The best run was chosen according to the highest log likelihood. For each run 10-fold cross-validation (CV) was computed.

### TreeMix analysis

Maximum likelihood tree construction and admixture modelling was performed with the TreeMix v. 1.12 software^39^ on the genome-based dataset of 36 populations and 64 individuals: on its original version or the version with CT and AG SNPs excluded. Initially, SNP window length (k) was optimized by testing various values (k=1, 5, 10, 20, 50, or 100) with 10 different random seeds, and the k setting producing the highest percentage of variance explained by the model was selected. Final runs were performed with the following settings: SNP window length, 5; number of migration edges from 0 to 8; trees rooted with the San population; global tree rearrangements used (option ‘-global’); no sample size correction (option ‘-noss’); 100 iterations, selecting a tree with the highest likelihood (and with the highest percentage of explained variance among trees with identical likelihoods). No pre-defined migration events were incorporated. One hundred bootstrap replicates re-sampling blocks of 5 SNPs were calculated and respective trees were constructed for 6, 7, and 8 migration edges using TreeMix. Bootstrap support values for nodes were mapped on the original trees using RAxML. Bootstrap values for migration edges were interpreted as described in Suppl. Information, Section 9, and in the respective figure legends (Fig. 3).

### f_3_ and f_4_, statistics

We used three and four population tests (*f_3_* and *f_4_*) developed by Patterson, et al.^38^ and implemented in programs *threepop* and *fourpop* of the TreeMix^39^ package. The source code in C^++^ was modified to enable multithreading and computing the statistic for all population combinations containing a given population or population pair. SNP window length settings used for computing standard errors of *f_3_* and *f_4_* statistics are shown for each dataset in Suppl. Table 1. Statistic *f_3_*(O; A, X_1_)^38^ measures relative amount of genetic drift shared between the test population A and a reference population X_1_, given an outgroup population O distant from both A and X_1_. Outgroup *f_3_* statistic is always positive, and its values can be interpreted only in the context of the reference dataset X. Statistic *f_4_*(X, O; A, B)^38^ tests whether A and B are equidistant from X, given a sufficiently distant outgroup O: in that case the statistic is close to zero. Otherwise, the statistic shows whether X is more closely related to A or to B, hence *f_4_* may be positive or negative. The statistical significance of *f_3_* and *f_4_* values is typically assessed using a Z-score: a statistic divided by its standard deviation. A threshold Z-score of 3 corresponds to a *p*-value of 0.00135 and is commonly used in case of multiple testing.

### Plotting

Plotting of various statistic values or sampling locations on the world map was performed using an open source software QGIS v.2.8 (http://qgis.org/en/site/) with open-source maps.

## Acknowledgements

We are grateful to all sample donors, and to local community members for their help in sample collection. We thank Eske Willerslev, Simon Rasmussen, Maanasa Raghavan, Iñigo Olalde, Andrés Ruiz-Linares, David Reich, Noah Rosenberg and Ripan Malhi for sharing genotyping and sequencing data. We would like to thank National Genographic and Family Tree DNA for genotyping our samples, Shi Yan (Fudan University, Shanghai, China) and Horolma Pamjav (Institute of Forensic Medicine, Budapest, Hungary) for their help with compiling Y-chromosome haplogroup frequency tables. Special thanks go to Alexey S. Kondrashov for putting our team together. P.F., P.C., and A.Z. were supported by the Moravian-Silesian region projects MSK2013-DT1, MSK2013-DT2, and MSK2014-DT1, and by the Institution Development Program of the University of Ostrava. T.V.T. was supported by grants from The National Institute for General Medical Studies (GM068968), the Eunice Kennedy Shriver National Institute of Child Health and Human Development (HD070996), and National Science Foundation Division of Evolutionary Biology (1456634). M.D.L. and M.S.G. were supported by the Russian Science Foundation: project nos. 14-50-00150 and 14-24-00155, respectively.

## Author contributions

P.F. and T.V.T. designed the study, took part in sample collection, performed data analyses, and wrote the paper. I.V.T., O.P.K. and T.N. were responsible for sample collection and manipulation, M.D.L. was responsible for genome sequencing, E.S.G. for software re-design, and A.Z., E.E.K., M.S.G., M.T., N.E.A., O.F., P.C., P.T., V.V.S. performed data analyses. G.S. contributed the linguistics section of the paper. Y.V.N. helped prepare the final version of the manuscript. All authors took part in interpretation and discussion of the results.

